# Mapping drinking water and quality of life aspects in urban settings: an application of the urban exposome

**DOI:** 10.1101/401927

**Authors:** Xanthi D. Andrianou, Chava van der Lek, Pantelis Charisiadis, Solomon loannou, Kalliopi N. Fotopoulou, Zoe Papapanagiotou, George Botsaris, Carjin Beumer, Konstantions C. Markris

## Abstract

Cities face rapid changes leading to increasing inequalities and emerging public health issues that require cost-effective interventions. The urban exposome framework constitutes a novel approach in tackling city-wide challenges, such as those of drinking water quality and quality of life. In this proof-of-concept study, we presented part of the urban exposome of Limassol (Cyprus) focusing on chemical and microbial drinking water quality parameters and their association with urban neighborhood indicators. A perceptions study and an urban population study was conducted. We mapped the water quality parameters and participants’ opinions on city life (i.e. neighborhood life, health care and green space access) using quarters (small administrative areas) as the reference unit of the city. In an exploratory environment-wide association study analysis, we used all variables (questionnaire responses and water quality metrics) to describe correlations between them accounting, also, for self-reported health status. Overall, urban drinking-water quality using conventional indicators of chemical (disinfection byproducts-trihalomethanes) and microbial (coliforms, E. coli, and Enterococci) quality did not raise particular concerns. The general health and chronic health status of the urban participants were significantly (FDR-corrected p-value<0.1) associated with different health conditions such as hypertension and asthma, or having financial issues in access to dental care. Additionally, correlations between trihalomethanes and participant characteristics (e.g. household cleaning, drinking water habits) were documented. This proof-of-concept study showed the potential of using integrative approaches to develop urban exposomic profiles and identifying within-city differentiated environmental and health indicators. The characterization of the urban exposome of Limassol will be expanded via the inclusion of biomonitoring tools and untargeted metabolomics platforms.

## 1. Introduction

The definition of the exposome in 2005 by Dr. Wild introduced a paradigm in environmental and population health research, promoting studies that either encompass simultaneous assessment of multiple exposures of the general population or focus on specific time windows of susceptibility (i.e. pregnancy), to capture the totality of environmental/lifestyle/behavioral exposures (1–4). Defining the exposome along with the advances in methodologies for high throughput analysis in shorter time has also redefined the study paradigm in environmental health. Thus, decoding the exposome will not benefit only environmental health, but it will lead to better understanding of disease development and progress. Additionally, it fosters innovation in exposure assessment, and it allows for intra-and inter-disciplinary approaches in public health to become more widespread than they are now. Within this context, more “exposomes” have been defined to address different totalities and with different units of reference (5–7).

Cities are dynamic and complex systems that will become the future focus of public health systems, because they currently host more than half of the global population and generate >80% of the global GDP (8, 9). Thus, cities are the main unit of reference for the urban exposome which can be seen in terms similar to the human exposome, as the totality of indicators (quantitative or qualitative) that shape up the quality of life and health of urban populations (10). In this context, the urban exposome, defined as the continuous monitoring of urban health indicators which refer to external and internal to the city parameters, is not merely the sum of individual exposures, but places cities in the center of an urban-oriented study framework, i.e. that of the urban exposome (10). In the urban exposome framework, the quality of life in urban centers is concurrently assessed along with other indicators, such as water quality, or prevalence/incidence of communicable and non-communicable diseases.

The value of clean water in public health is historically acknowledged at all societal levels. As more people nowadays live in cities, easy access to safe and affordable water is becoming more important in eliminating possible within-city health and societal disparities.

Besides the technical provisions to maintain water of good quality, the uninterrupted availability of water becomes an issue due to climate change manifestations, especially in areas that are now or expected to be hit harder by extended droughts and other related meteorological phenomena. Europe, overall, is expected to face increases in both the extent of geographical areas affected by droughts and in the duration of such climatic events (11). Within this context, cities located in the Southeast Europe and the Mediterranean region are predicted to face challenges in maintaining ample water availability and excellent water quality in the future (11).

The biological and societal networks in an urban setting that emerge from various aspects of climate change, water availability, water quality and population health are becoming increasingly complex to solely rely upon classical urban health approaches. Cities and their smaller areas (i.e. neighborhoods, or small areas) are warranted to address water-related issues, such as water demand, safety, security and quality issues, while tackling societal inequalities and health disparities. In this context, the concept of the urban exposome is introduced to help scientists and policy makers to better articulate the framework of the systematic spatio-temporal surveillance and monitoring of a city’s heterogeneous health profile. The urban exposome is central to better understanding urban health dynamics and goes beyond the human exposome framework, which is instrumental in personal health assessment or identifying risk factors for different health outcomes in population studies. The aim of this study was to map different indicators of the urban exposome that mostly related to drinking water and quality of life as part of a larger study that was set up a case study to present the application of the urban exposome framework in the city of Limassol, Cyprus.

Limassol is a coastal city, the biggest port of Cyprus, and it is defined as a medium-sized city (~200000 inhabitants according to the 2011 Population Census of Cyprus) (12, 13), currently facing rapid economic development. Half of the urban population of the urban area of Limassol resides within the municipality of Limassol (~110000 inhabitants) in the center of the city. The objective of this study was to describe the water quality and aspects of quality of life following the integrative urban exposome framework in the municipality of Limassol, having quarters (small within-municipality administrative areas) as the unit of reference. This urban study was *a priori* designed without a hypothesis-testing, or an one exposure-one outcome approach, because of the inherent nature of the urban exposome framework that relies upon a hypothesis-generating scope, instead. Our intention was to provide a practical example and a proof-of-concept study on how the urban exposome framework could be used to study drinking water quality aspects in a dynamic urban setting of a typical mid-sized city.

## 2. Materials and Methods

### 2.1. Application of the urban exposome in the municipality of Limassol, Cyprus

We have previously described the urban exposome as all indicators that need to be continuously monitored for the assessment of city health (10). The urban exposome focuses on cities and their geographic (sub)areas as the study units and it parallels the human exposome characteristics (10). Within the framework of the urban exposome, external to the city parameters that cannot be influenced by the city itself can be either general (e.g. global trends and policy decisions) or more specific (e.g. climate change impacts, demographic changes, culture). It follows that internal parameters are those that are integral to the city, such as infrastructure, built/neighborhood environment and determinants of population health (e.g. socioeconomic factors). An exposome-based study was conducted in summer to describe the water quality aspects of the urban exposome of Limassol, following an integrative and interdisciplinary approach (Figure 1). This approach included the following parts:

- A perceptions survey with a mixed-method approach, which was conducted to evaluate the perceptions of stakeholders (i.e. citizens and municipality officials) about the quality of life and certain environmental risks (e.g. drinking water chemical and microbial risks) in the city.
- A cross-sectional urban population study with a short questionnaire and collection of tap water samples from households distributed in the quarters of the municipality of Limassol to evaluate water quality indicators and citizen’s attitudes on the environment, quality of life in the city and their health status.

The comprehensive assessment of the urban exposome for the Limassol city goes beyond the scope of the present analysis that focuses on water quality and perceptions about life in the city.

**Figure 1.**
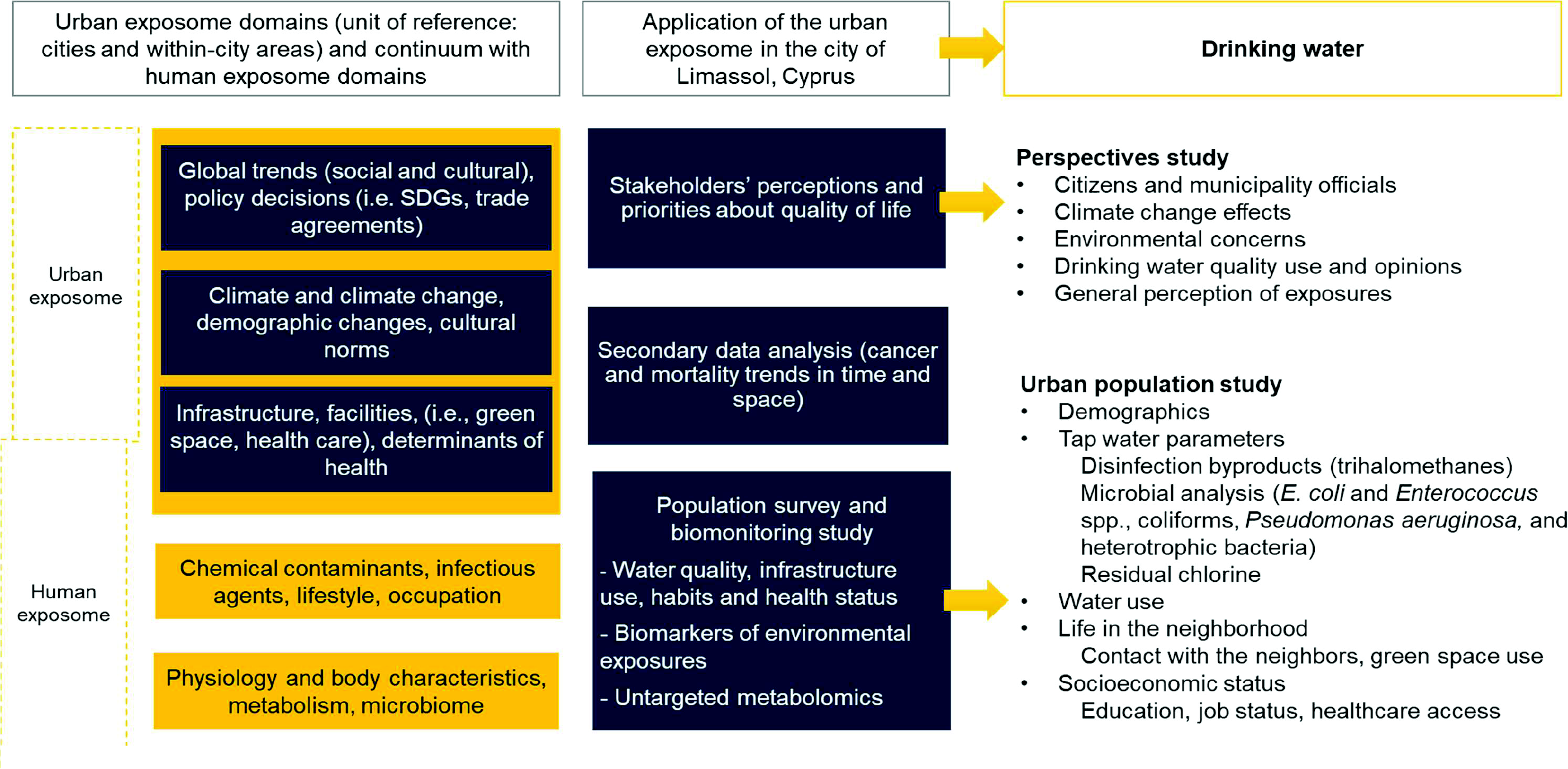
Urban exposome - human exposome continuum, and the practical application of the urban exposome framework in the urban setting of Limassol city. The parts specifically discussed in the current analysis include a perceptions study and an urban populationstudy, which includes parameters measured in drinking-water to assess the water quality coupled with questionnaire responses about individual lifestyle, behavioral, and personal health indicators

### 2.2. Perceptions study

The perceptions study was an urban community-based participatory research to actively engage the municipality of Limassol officers and its citizens in urban health issues which were identified as the community stakeholders (14). The stakeholders’ perceptions were assessed using a mixed methods approach. With the municipality functionaries (i.e. technical officers from the municipality of Limassol and neighboring municipalities) face-to-face interviews were conducted, and short questionnaires were administered. For the citizens’ perceptions, an online anonymous questionnaire initially distributed via email among staff at the Cyprus University of Technology campus that is located in the municipality of Limassol, followed by its distribution to the general public via mailing lists and social media (Facebook). The questions asked during the face-to-face interviews and the questionnaire for the municipality officers focused on the identification of trends shaping the city of Limassol, assessment of the climate change manifestations and their impact, scoring of environmental health concerns, an assessment of what was believed to be the citizens major health threats with regards to urban life, and perspectives about future opportunities to improve health in the city. The questionnaire administered to the citizens included various questions on climate change perceptions, environmental concerns in general, health perceptions and perceptions about drinking-water quality. The descriptive analysis of the questionnaire responses (number of respondents and frequencies) was conducted in SPSS 20 (IBM Corp. Armonk, NY: IBM Corp.).

### 2.3. Urban population study

A cross-sectional population study was set up in the municipality of Limassol, Cyprus. Participants were randomly recruited in summer 2017 from all quarters of the municipality following the 2011 census population distribution (Table S 1). For this study, we also collected urine samples for biomonitoring purposes (not included in this analysis). Thus, sample size estimations were based on the assumption that a total of 120 participants randomly selected from the whole municipality would allow us to evaluate the baseline levels of environmental exposures, since according to previous biomonitoring studies a sample of at least 120 randomly selected participants is adequate to capture the 95^th^ percentile of the population levels (15). In the analysis, two small in area/population size quarters with one participant were merged with neighboring ones and three quarters located along the beachfront, each having one participant were also merged together (Figure S 1).

Tap water samples were collected from participating urban households and *in situ* measurements of free chlorine were taken during house visits. All participants were asked to fill in a questionnaire that included, among others, questions about life in their neighborhoods, self-reported health status and drinking water habits. The questionnaire was based on previous studies on urban health and the European Health Survey questionnaire (16, 17). The study was approved by the National Bioethics Committee of Cyprus (decision number: 2017/23).

### 2.4. Water sampling and analysis

The tap water faucet was externally cleaned with ethanol prior to sample collection, and the water was then left to flow freely for ~30 seconds. Tap water samples for trihalomethanes (THM) analysis were collected in falcons containing oxidation preservative mixture, while the water samples used in the microbial analysis were collected in sterile polypropylene vials. THM analysis was conducted according to the previously published methods by Charisiadis et al. (18). All four THM species were measured in the collected tap water samples from the participating households: chloroform (TCM), bromodichloromethane (BDCM), dibromochloromethane (DBCM), and bromoform (TBM). The limits of detection (LOD) were 0.13μg/L for TCM and DBCM, while it was 0.11μg/L for BDCM and TBM. The microbial analysis was conducted after the water was cultured onto selective media for the detection and enumeration of total coliforms, *E. coli, Enterococcus* spp., *Pseudomonas aeruginosa*, and total viable counts (TVC) at 22 °C and 37 °C. The methods used for the microbial analysis are presented in Supplementary Information Section A.

All water samples were collected from the main faucet of the household used to satisfy their potable needs and it was directly connected to the municipality’s water supply. In case a point of use market filter was present, it was removed to collect unfiltered tap water; for those households (n=13) having point of use filters that were permanently connected to the faucet, tap water samples were excluded from the main statistical analysis.

Residual chlorine was measured with a portable photometer (MaxiDirect, Lovibond) in water directly collected from the tap using the DPD (N,N-diethyl-p-phenylenediamine) method.

### 2.5. Statistical analysis for the population study

#### 2.5.1. Descriptive analysis of the questionnaire responses

Descriptive statistics (i.e. means and standard deviation for the continuous variables, and frequencies and percentage by category for the categorical variables) were calculated for the responses to the questionnaire. The descriptives were grouped by category of question (i.e. demographics and other background characteristics, drinking water habits and cleaning activities, lifestyle and behavioral indicators, healthcare services access, health status, life in the neighborhood) for the complete study population (n=132).

#### 2.5.2. Descriptive analysis of the water indicators and water habits

Descriptive statistics for the drinking water THM and microbial counts were separately presented for the samples collected directly from the tap when no filter was attached (n=119) than those 13 samples that were collected with a filter present. When THM values were below the LOD (n=4 for TCM and n=1 for TBM), they were imputed to LOD/2. The sum of all THM species (total THM) and the sum of the brominated species (i.e. BDCM, DBCM, and TBM; the BrTHM) were calculated. For the statistical analysis of the microbial water quality, we considered the presence or absence of colonies for *E. coli* and *Enterococcus spp*. instead of the absolute count number (19). For the coliforms, *Pseudomonas aeruginosa*, and TVC at 22 and 37 °C the presence or absence the percentage of samples with detected colonies were described. Specifically, for TVC at 22 and 37 °C a smaller number of samples was analyzed (n=95) due to external contamination. For the THM concentrations the descriptives included: mean, standard deviation (sd), median and percentiles, i.e. 25^th^, 95^th^ percentiles, and the range, i.e. min and max, while for the microbial analysis the frequency of samples with at least one CFU (and the percentage) was calculated.

The results of the THM and microbial analyses were presented separately for the total THM, the *E. coli* and the *Enterococci* spp., as they are considered of higher priority compared to the single THM species and the other microbial indicators (e.g., coliforms).

#### 2.5.3. Mapping of the water and the quality of life indicators

To evaluate how indicators pertaining to the quality of life and water quality are distributed in different quarters of Limassol, we mapped a selection of urban indicators by quarter. For the water quality indicators, the median levels of total THM, free chlorine, BrTHM per quarter were mapped, as well as, the percentage of samples with detectable levels of coliforms and TVC. From the urban questionnaire data, all indicators were mapped. However, for brevity, we present and discuss one indicator per category of the following: (i) issues on access to health care services due to delays and financial constraints, (ii) life in the neighborhood, and iii) two indicators from the category on access to green spaces to illustrate potential differences between quarters. While all the indicators presented in the results for brevity maps were prepared for the ones on which the participants had most consensus within a category.

#### 2.5.4. Exploratory environment-wide association study (EWAS)

We used an exploratory environment-wide association study (EWAS) approach as part of the urban exposome framework to agnostically synthesize knowledge and concurrently investigate possible associations of the measured water quality indicators, questionnaire-based socioeconomic, lifestyle and behavioral factors, with three self-reported health status outcomes: (i) general health status, (ii) diagnosed with any chronic disease, and (iii) diagnosed with any disease in the past year (“any disease the past year”, i.e. asthma, diabetes, allergies, hypertension, cardiovascular or respiratory illnesses, depression, cancer, musculoskeletal problems) (20). The EWAS mode of analysis included a correlation matrix between all variables, followed by regressions, and a multi-omics-based approach where investigated correlations accounted for specific health outcomes. Below we describe the variables used in the EWAS and the details of the analysis.

Besides the three self-reported health outcomes, the variables included in the EWAS approach were divided in the following “blocks/groups”:

- Block/group 1: water THM levels, free chlorine
- Block/group 2: drinking water habits (e.g., number of glasses of water consumed by source)
- Block/group 3: household cleaning activities
- Block/group 4: neighborhood quality of life (variables of “highest consensus”: heath care access, life in the neighborhood and green urban spaces)
- Block/group 5: participant characteristics (e.g., age, sex, BMI)
- Block/group 6: self-reported diseases in the past year (e.g. asthma, diabetes, allergies, hypertension, cardiovascular or respiratory illnesses, depression, cancer, or musculoskeletal problems).

In all categorical variables the “I don’t know/l don’t want to answer” responses were recoded to missing. Then, scores were added per category (presented in Table S 2) and used in statistical analysis when categorical variables could not be used (e.g. correlations).

In the preliminary correlation analysis, all variables (including the three outcomes) were used as continuous and the Spearman correlation coefficient was calculated without any transformations. The results were visualized with a correlation plot. Then, regression models were fitted. The variable for the general health status was used as continuous in linear regression whereas responses for selfreported chronic disease and any disease the past year were used as binary variables in logistic regression. The regression models were repeated after adjusting for age and sex. The continuous predictors were scaled and centered in all regression models. The p-values of all model parameters that were used for inference (i.e. excluding the intercept for the univariable models and excluding the intercept, sex, and age coefficients from the adjusted models) were summarized and corrected for Benjamini-Hoechberg false discovery rate (FDR). Only parameters with an FDR-corrected p-value <0.10 separately applied to the univariate (n=129 tests) and the adjusted models (n=123 tests) were considered statistically significant.

In the last part of the EWAS analysis, we followed an approach used in multi-omics studies where the variables are grouped and partial least squared discriminatory analysis (block PLS-DA) is conducted to identify possible correlations between the variables of the different blocks accounting for an outcome (21). In this analysis, the predictor variables were used as continuous and all outcomes were used as categorical. The correlations between the predictor variables in blocks were presented in circular plots (circo plots) where positive and negative Pearson pairwise connections are shown in the circle and lines indicate the levels of each variable within each outcome category.

All analyses (regressions and block PLS-DA) were performed for the three outcomes using all the blocks/groups as predictors except for the analysis for the outcome “any disease the past year”. This variable was created as the summary of the variables of block/group 6 (self-reported diseases the past year). Thus, in this analysis, the separate diseases of block/group 6 were not included due to their association with the outcome of “any disease the past year”.

All analyses were conducted in R 3.5.1 with RStudio 1.1.423 (22, 23). The data and the scripts used in the analysis can become available upon request. The packages used in the analysis are listed in Supplementary Information (R packages used in the data analysis).

## 3. Results

### 3.1. Perceptions study results

In the face-to-face interviews and through the short questionnaires the importance of climate change and its effects was pointed out by all municipality technical officers (n=6). In rating the environmental concerns (1 for very low and 5 for very high concern), water quality had the lowest score with increasing order of scoring for soil contamination, waste, general chemical exposures, noise and air pollution.

Out of 134 questionnaire respondents from the general public, 91 reported living in Limassol (35% males and 65% females) with a mean age of 35 years old [range:18-77 years old]. A large majority of them (81%) was born on Cyprus, and 13% was born in another EU country, the rest (6%) were born in a non-EU country. Most of the respondents (84%) were highly educated holding at least a Bachelor’s degree. Around half of them were married (46%) with children (47%).

Residents of urban Limassol were mostly concerned about being severely exposed to air pollution and noise (Table 1). Water quality ranked low while air pollution and noise ranked high in the severely exposed category among all environmental exposures. Approximately 30% of respondents reported that they were not exposed to water pollution or soil contamination, but at the same time an equally high proportion of respondents reported “don’t know”, suggesting inadequate knowledge about the drinking-water or soil quality in their city. With regards to water quality, 81% reported worrying about chemical exposures, and 41% reported they were exposed to chemicals on a daily basis.

**Table 1.** Perceptions about environmental exposures among the respondents from the city of Limassol (n=91).

|  | **Severely exposed** | **Somewhat exposed** | **Not exposed** | **Don't know** |
| --- | --- | --- | --- | --- |
| Noise (traffic, airplanes, factories, neighbours, animals, restaurants/bars/clubs) | 38 (42%) | 48 (53%) | 5 (5%) | 0 (0%) |
| Air pollution (fine dust, grime, fume, ozone) | 40 (44%) | 40 (44%) | 10 (11%) | 1 (1%) |
| Bad smells (industry, agriculture, waste) | 15 (17%) | 35 (38%) | 39 (43%) | 2 (2%) |
| Water pollution (microbes/chemicals in drinking water) | 8 (9%) | 22 (24%) | 31 (34%) | 30 (33%) |
| Soil contamination (eg., chemical waste dump) | 5 (5%) | 15 (17%) | 32 (35%) | 39 (43%) |

The following information from the analysis of the questionnaires answered by the residents of Limassol also related to the use of water and concerns/behaviors: among all urban respondents, only 29% reported drinking the water straight from the tap, while 32% reported treating the water before consumption (filtering or cooking), and 39% mentioned not drinking the tap water at all. Also, among the highest in importance concerns of the citizens about tap water quality were those associated with either chemicals (47%), or microbes (37%) and much less concerned about the taste (10%).

### 3.2. Population study results

#### 3.2.1. Background information and opinions

In total, 132 residents of the Limassol municipality answered the questionnaire and agreed to water collection from their household’s main tap. The distribution of the study participants by quarter and the population can be found in Table S 1. The mean age was 45.6 years and the majority were females (62.1%). Most of the participants were born in Cyprus [n=114 (86.4%)] living there for all their lives (Table 2, Table S 3).

**Table 2.** Background characteristics of the 132 participants of the urban population study conducted in Limassol, Cyprus (2017).

|  | **Overall (n=132)** |
| --- | --- |
| **Age (mean (sd))** | 45.6 (13.2) |
| **Sex (%)** |  |
| Females | 82 (62.1) |
| Males | 50 (37.1) |
| **BMI (mean (sd))** | 26.25 (5.06) |
| **Marital Status (%)** |  |
| Cohabiting | 8 (6.1) |
| Divorced | 15 (11.4) |
| Married | 82 (62.1) |
| Single | 26 (19.7) |
| Widowed | 1 (0.8) |
| **Education (%)** |  |
| Primary school | 2 (1.5) |
| Gymnasium | 2 (1.5) |
| High School/Technical School | 28 (21.4) |
| Non-university tertiary education | 16 (12.2) |
| Post-secondary education | 6 (4.6) |
| Bachelor's degree (BSc/BA) | 45 (34.4) |
| Master's degree or doctorate (MSc/MA, PhD) | 32 (24.5) |

Most of the study participants reported being in very good or good health condition (46% and 43%, respectively). However, 21% reported having a chronic disease and 57% reported at least one of the following health conditions during the past year: asthma, cardiovascular diseases, hypertension, diabetes, liver conditions, cancer, depression, or musculoskeletal problems (Table 3). With regards to access to health care, the main questions were about delays due to lack of transportation or long waiting lists (Table S 4). Lack of transportation did not seem to be a major constrain to access health care centers among those that opted to answer; however, long waiting lists were reported by 11 %. In the question about financial constraints for health care access, delays in dental care were most frequently mentioned (14%). Access to urban green space was easy since most participants (64%) reported living close to green spaces, but a total of 63% also reported that these green spaces were not well-maintained and there was a consensus not using them. A summary of the responses about health care access, lifestyle, the quality of life in the neighborhood and other urban topics can be found in Table S 4.

**Table 3.** Health status indicators assessed through the questionnaire among the 132 participants of the urban population study (Limassol, Cyprus; 2017).

|  |  | Overall (n=132) |
| --- | --- | --- |
| **General health assessment (%)** | Very good | 61 (46.2) |
|  | Good | 57 (43.2) |
|  | So and so | 12 (9.1) |
|  | Bad | 1 (0.8) |
|  | Very bad | 1 (0.8) |
| **Chronic disease (%)** | Do not know | 5 (3.8) |
|  | I don't want to answer | 3 (2.3) |
|  | No | 97 (73.5) |
|  | Yes | 27 (20.5) |
| **Any disease in the past year (%)** | No | 57 (43.2) |
|  | Yes | 75 (56.8) |

#### 3.2.2. Water quality indicators assessment of drinking water habits

The main chemical water quality indicators assessed were the THM. Only 2% of the households’ tap water exceeded the THM parametric value (100μg/L). Results conforming with the parametric values were also obtained for the microbial indicators monitored, i.e. *E.coli* and *Enterococci* spp. All samples were within the parametric values (0 CFU per 100mL), besides one household where Enterococci colonies were detected. Total coliforms were detected in 28 of the 132 households and *Pseudomonas aeruginosa* counts were detected in 5 out of the 132 households (Tables 4 and 5).

**Table 4.**
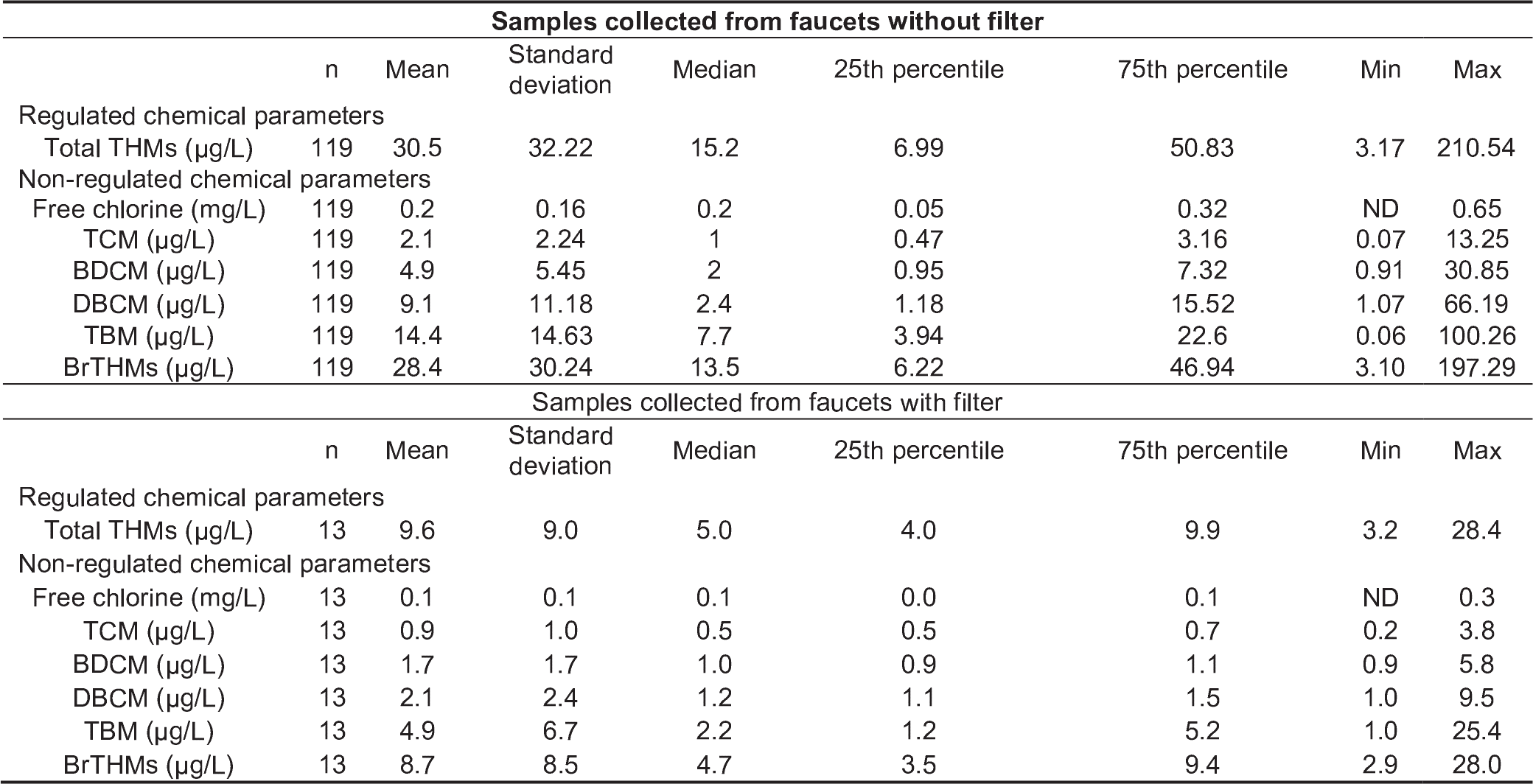
Chemical drinking water parameters analyzed in water samples collected in Limassol, Cyprus (2017) for n=119 samples collected from faucets without point of use filter and for n=13 samples collected from faucets with a point of use filter that could not be removed.

**Table 5.**
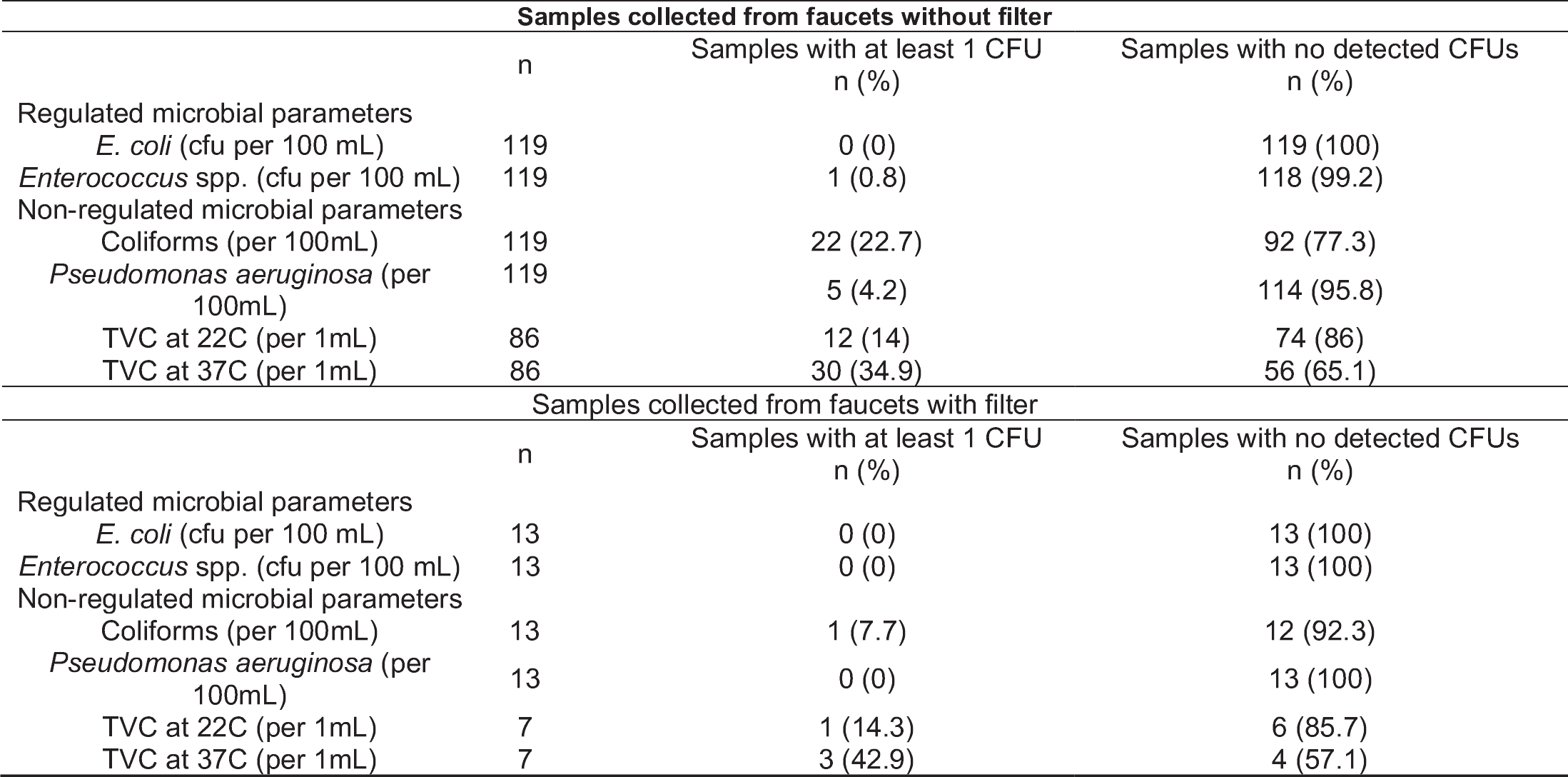
Microbial drinking water parameters analyzed in water samples collected in Limassol, Cyprus (2017) for n=119 samples collected from faucets without point of use filter and for n=13 samples collected from faucets with a point of use filter that could not be removed.

| Samples collected from faucets without filter |  |  |  |
| --- | --- | --- | --- |
|  | n | Samples with at least 1 CFU<br>n (%) | Samples with no detected CFUs<br>n (%) |
| Regulated microbial parameters |  |  |  |
| *E. coli* (cfu per 100 mL) | 119 | 0 (0) | 119 (100) |
| *Enterococcus* spp. (cfu per 100 mL) | 119 | 1 (0.8) | 118 (99.2) |
| Non-regulated microbial parameters |  |  |  |
| Coliforms (per 100mL) | 119 | 22 (22.7) | 92 (77.3) |
| *Pseudomonas aeruginosa* (per 100mL) | 119 | 5 (4.2) | 114 (95.8) |
| TVC at 22C (per 1mL) | 86 | 12 (14) | 74 (86) |
| TVC at 37C (per 1mL) | 86 | 30 (34.9) | 56 (65.1) |
| Samples collected from faucets with filter |  |  |  |
|  | n | Samples with at least 1 CFU<br>n (%) | Samples with no detected CFUs<br>n (%) |
| Regulated microbial parameters |  |  |  |
| *E. coli* (cfu per 100 mL) | 13 | 0 (0) | 13 (100) |
| *Enterococcus* spp. (cfu per 100 mL) | 13 | 0 (0) | 13 (100) |
| Non-regulated microbial parameters |  |  |  |
| Coliforms (per 100mL) | 13 | 1 (7.7) | 12 (92.3) |
| *Pseudomonas aeruginosa* (per 100mL) | 13 | 0 (0) | 13 (100) |
| TVC at 22C (per 1mL) | 7 | 1 (14.3) | 6 (85.7) |
| TVC at 37C (per 1mL) | 7 | 3 (42.9) | 4 (57.1) |

The drinking-water consumption habits reported by the urban participants similarly reflected what was already observed in the perceptions study (presented earlier) where only 30% reported consuming tap water and most participants reported a preference to other sources, such as bottled water (Table 6). Among the participants of this part of the study, the majority reported using tap water in general, however bottled water use was noted by most respondents. The more frequently reported single drinking water source was bottled water (30%) followed by tap water (22%). A comparable proportion of the participants reported the combined use of tap water and bottled water (Table 6). More than half of the study partipants reporting consuming less than one glass of water per day from the tap (median number of glasses consumed from tap was 1 glass/day) (Table 6).

**Table 6.**
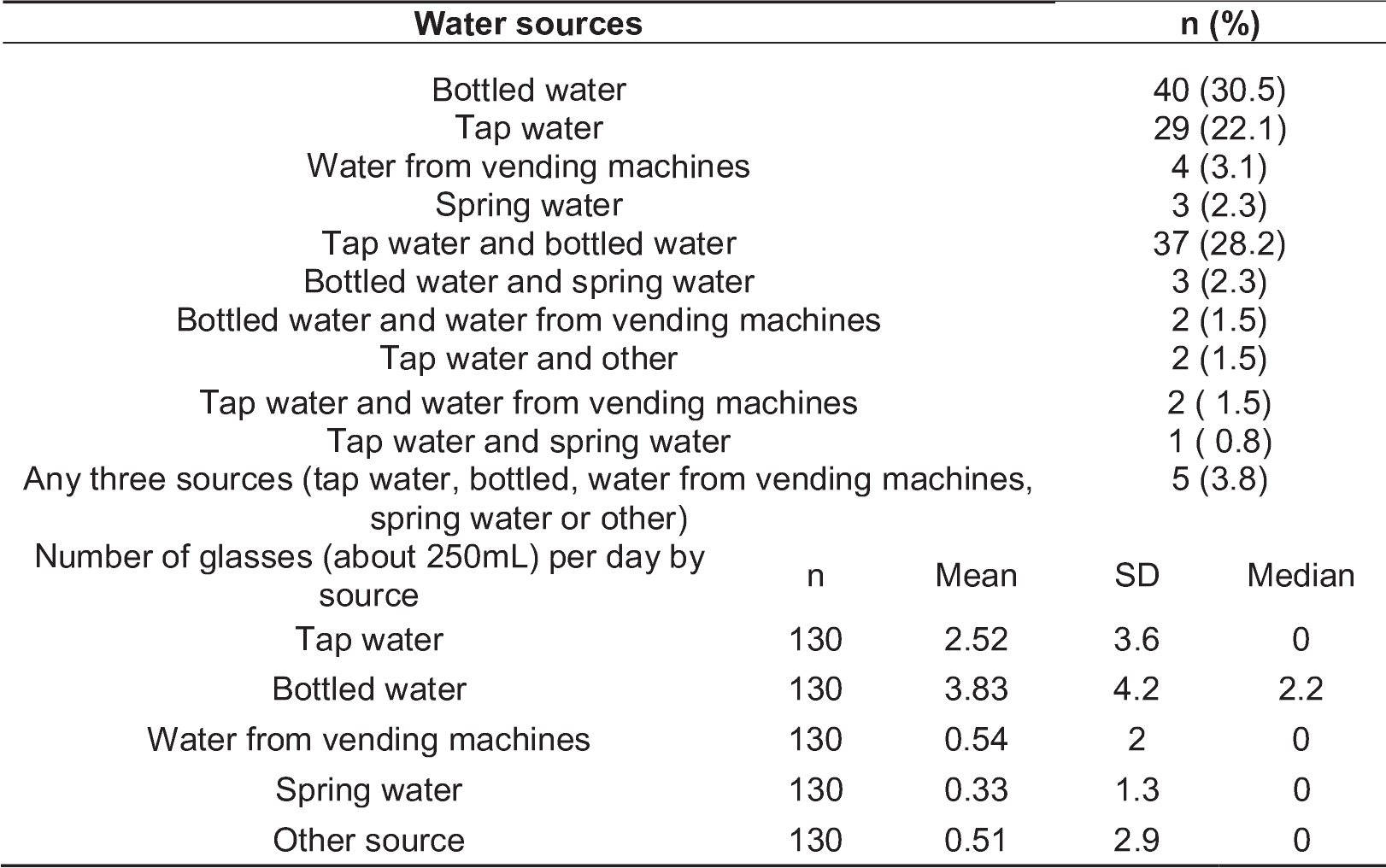
Self-reported choices of water sources by the study participants (n=132) of the urban population study in Limassol, Cyprus (2017).

### 3.3. Mapping of urban indicators of water and quality of life

The measured water quality indicators in each household were aggregated and mapped by quarter. For the chemical water parameters, the mapped median total THM and the brominated species (BrTHM) by quarter showed similar patterns (Figure 2). As the BrTHM are a subset of the total THM, their median values by quarter were lower. The highest median THM values were observed in the quarters located between the beachfront and the northern quarter of Agia Fylaxi (near the center of the city). Mapping of free chlorine levels followed an opposite pattern to THM, i.e. higher levels of free chlorine in the seaside quarters and slightly higher in the northern quarter (being closer to the main water treatment plant). The maximum median value of total THM was 63μg/L and it was observed in the center of Limassol (quarter of Agios Georgios). In this quarter, the median free chlorine levels were below detection. The range of the median total THMs by quarter was 6-63μg/L for the small quarters in the beachfront and behind the city port and the quarter of Agios Georgios. The variability of the median free chlorine levels was smaller, ranging from below detection to 0.4mg/L in the quarters of Agia Zoni and Agia Trias (in the center and by the seafront, respectively). A map with the quarter names of municipality of Limassol is available in Supplementary Information (Figure S 1).

With regards to the microbial parameters, as mentioned in the previous section, only one sample had detectable colonies of *Enterococcus spp*. while *E. coli* was not detected at all. Thus, we mapped only the percentage of samples found positive for coliforms or had detectable heterotrophic bacteria (TVC at 22 and 37C). In a quarter, two of the three samples analyzed were positive for coliforms and, therefore, it had the highest percentage of samples with colonies. This quarter was geographically located in the zone with the highest median chlorine levels (0.3 mg/L) (Figure 2).

**Figure 2.**
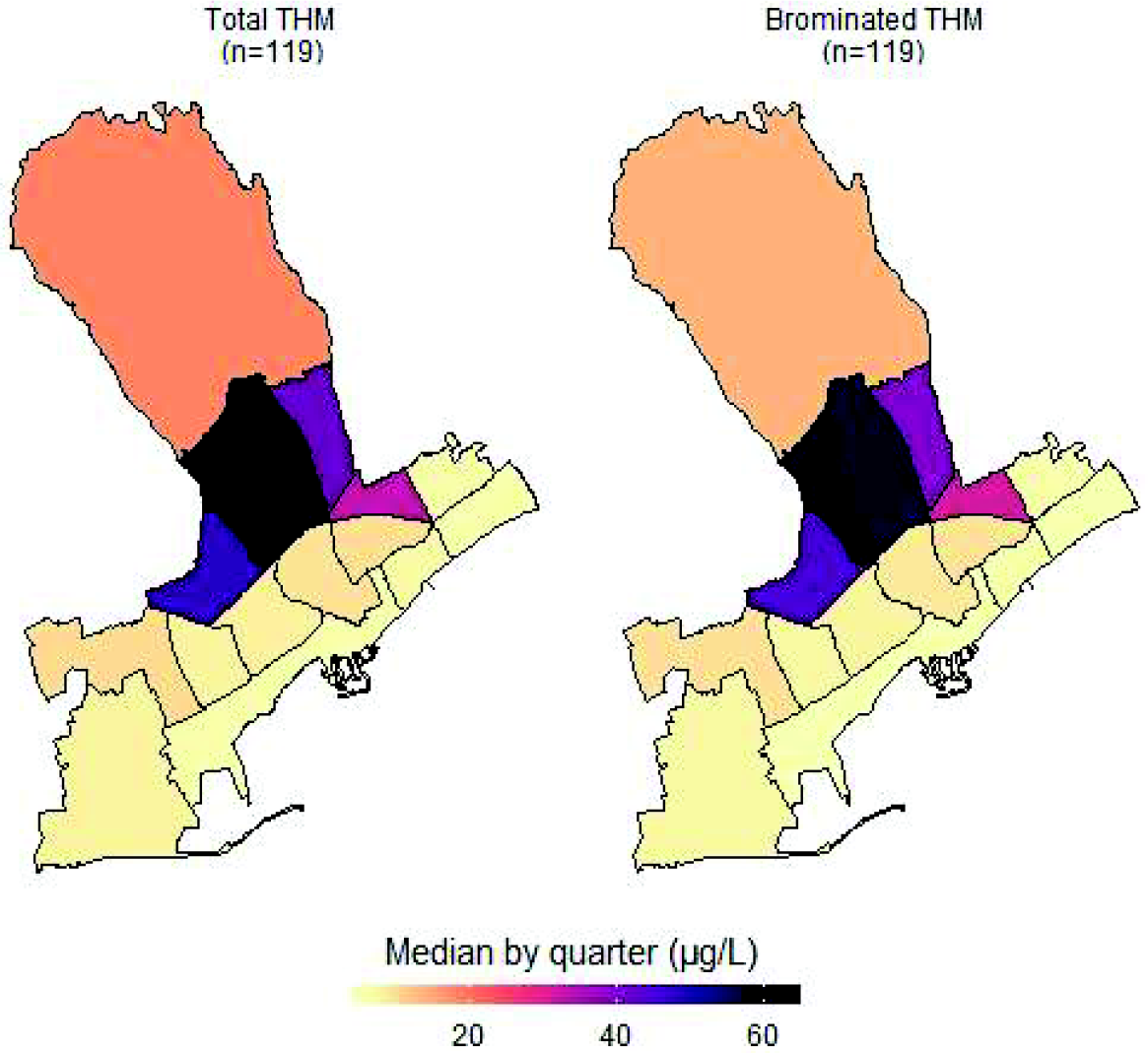
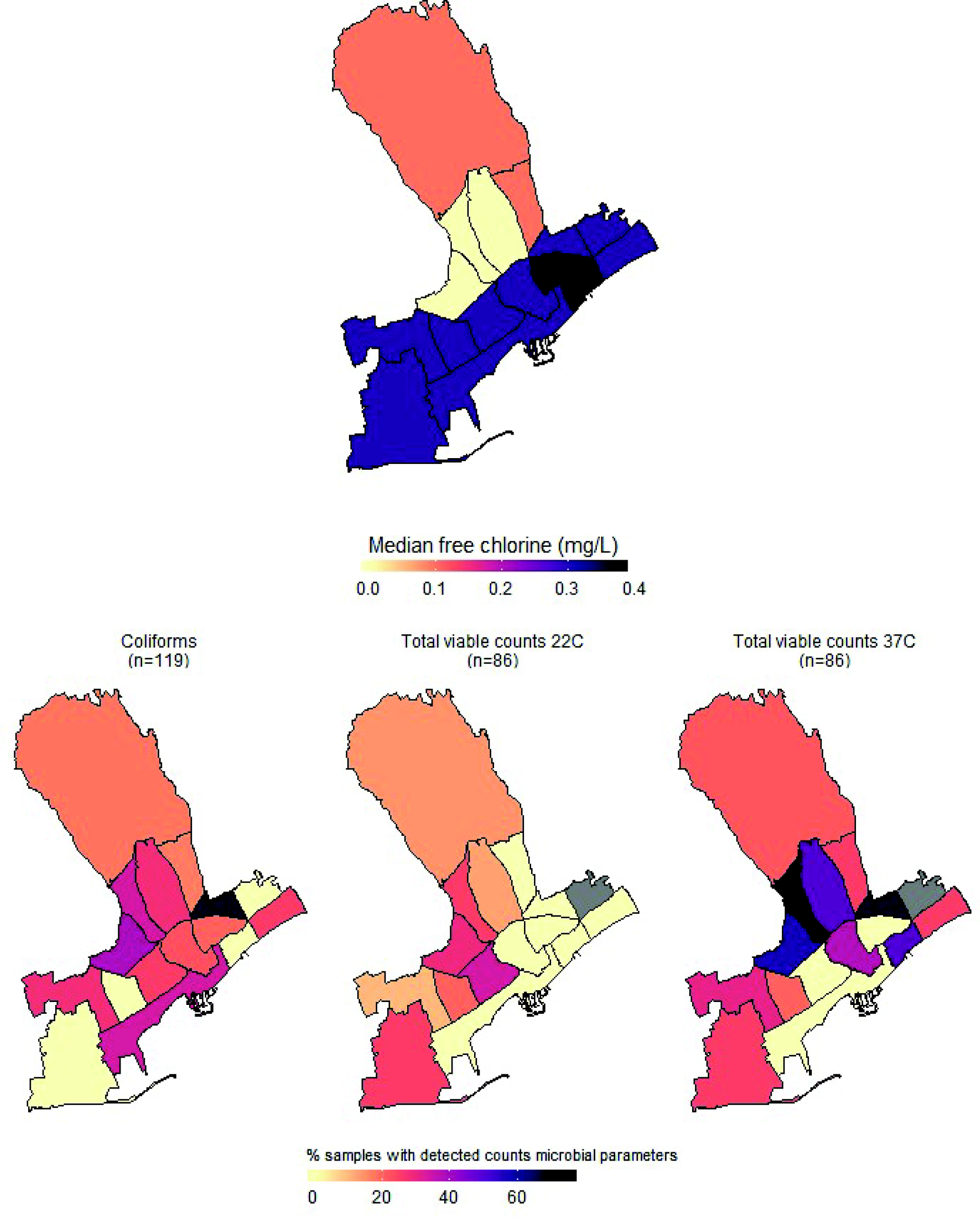
Maps of the median water total THM, BrTHM and free chlorine and of the percentage of samples with detectable counts of the monitored microbial parameters by quarter within the municipality of Limassol, Cypris (2017).

#### 3.2.3. Description and mapping of access to health care services, life in the neighborhood and use of green spaces

With regards to access to health care, most participants reported having issues with two major parameters, i.e., long waiting lists and financial constraints to access dental care. From the respective maps (Figure 3), it was evident that participants living in the quarters of Katholiki, Agia Trias, Omonoia and Agios Nektarios did not report any issues pertaining to access to health care. Whereas, other quarters such as Agios loannis/Arnaoutogeitonia, Agia Zoni and Agios Nikolaos were more consistently in the “mid-range” with 20% participants reporting issues for both indicators. With regards to the question about having someone in the neighborhood to ask for help in emergencies, overall, only 25/132 participants opted for the answers “I don’t know” and “I disagree completely or probably”. However, responses of strong agreement (“I completely agree” vs all the other options from “I probably agree” to “I completely disagree”) varied a lot across the quarters. For example, in Agios Spyridonas only 20% agreed that there is always someone to help them while in Agios Nikolaos 80% (Figure 2).

**Figure 3.**
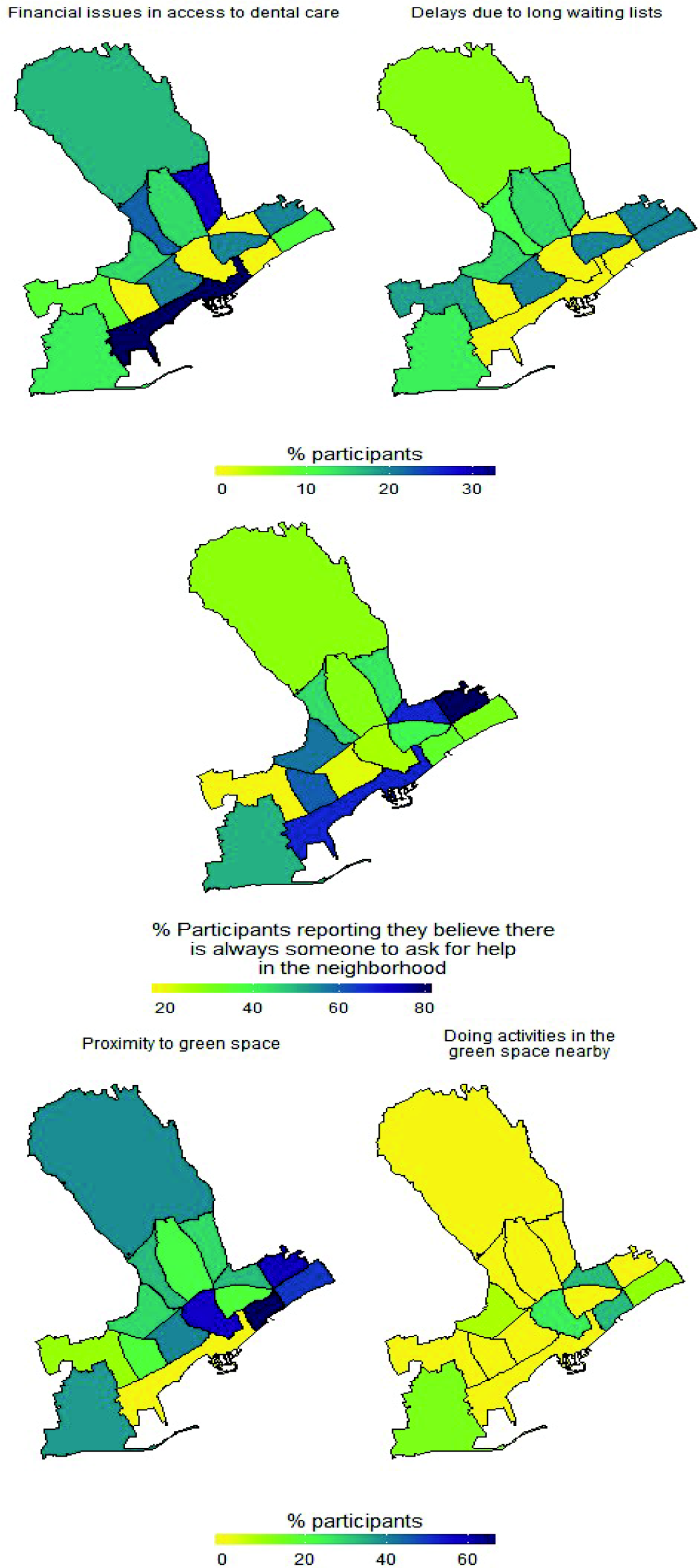
Maps by quarter of the percentage of study participants within the quarters of Limassol municipality, Cyprus (2017) agreeing with different statements about life in the neighborhood or access to green spaces.

### 3.4. Exploring environment-wide associations within the municipality of Limassol

A correlation plot between all variables used in the EWAS analysis (listed in Table S 2) did not show any specific or unexpected patterns of correlation among the urban variables (Figure 4). All THM compounds in drinking water correlated well with each other and they were negatively correlated with the free chlorine levels. Notable correlations were observed among different variables of the same block/group, besides for THM; for example, strong correlation was observed between household cleaning variables (mopping, dishwashing and bathroom cleaning) and certain health conditions, such as musculoskeletal problems (neck problems).

**Figure 4.**
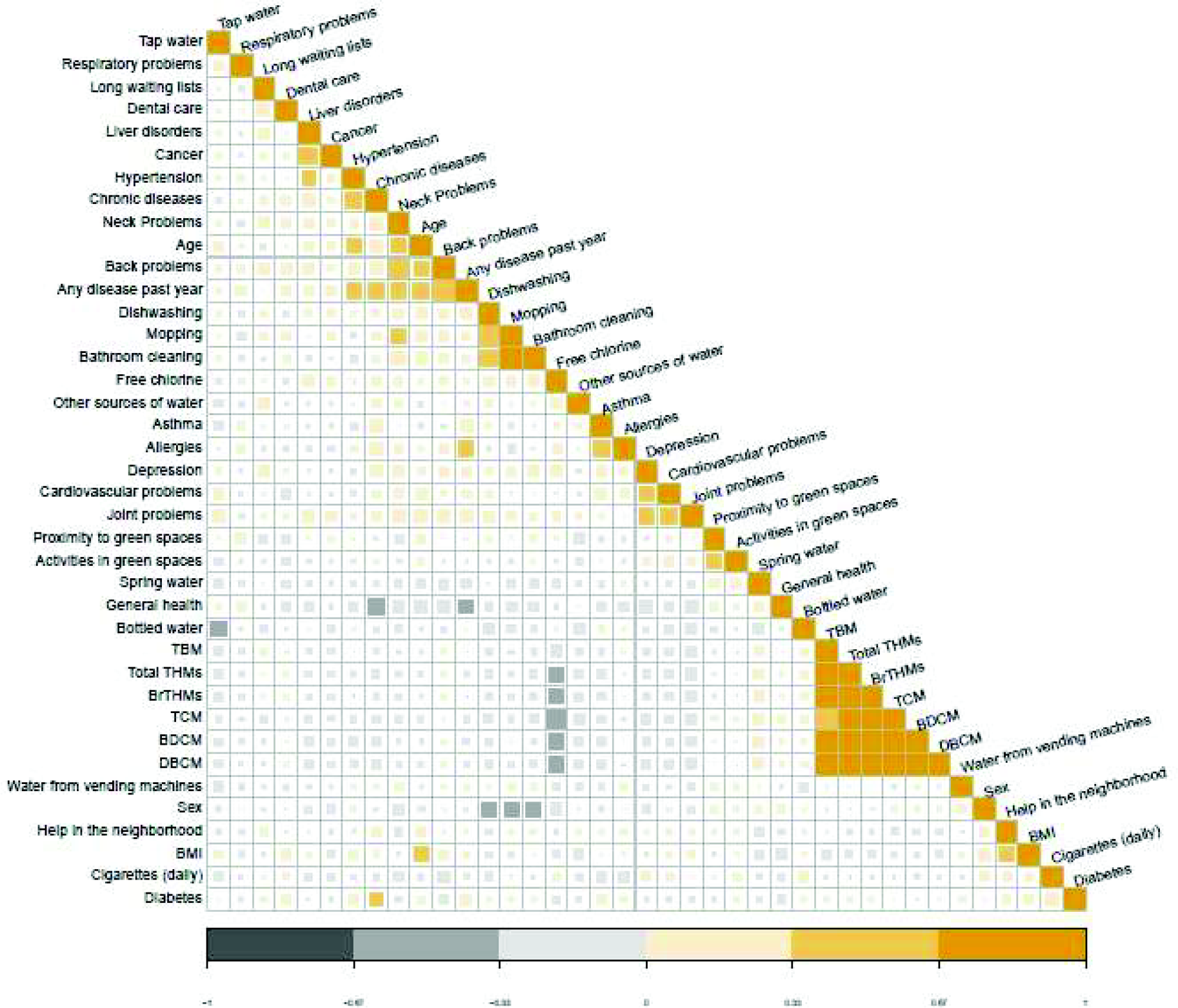
Correlation plot (Spearman correlation coefficient) for all the variables used in the environment-wide.

In the regression part of the EWAS analysis, a total of 129 predictors were summarized from the simple models and 123 predictors used in the models adjusted for age and sex. These predictors include both parameters measured in water, i.e. THMs, and questionnaire responses (summarized in Table S 2). In the models adjusted for age and sex, four parameters: i.e. financial issues in access to dental care, depression and hypertension and asthma had an FDR-corrected p-value <0.1. Having encountered financial issues in access to dental care [n=18 (14.1%) participants] and depression [n=3 (2.8%) participants) were statistically significant negative predictors for better general health status, while and higher odds of having a chronic disease were associated with hypertension and asthma [n=19 (16.2%) and n=10 (8.5%) participants, respectively]. In the univariate regression, in addition to the parameters that were significant in the adjusted models, musculoskeletal problems (i.e. neck and back problem) were associated with higher odds of having a chronic disease but not with the outcome of general health status. The first 20 parameters ranked by the FDR adjusted p-value can be found in Supplementary Information (Table S 5).

The results of the second EWAS analysis and the correlation between the variables used in the PLS-DA models were summarized in the circular plots (circo plots) of Figure 5. Again, all three outcomes were used, however, the model for the chronic disease did not converge and thus, the plot is not presented. For the other two outcomes the associations between the variables were not the same, as expected due to the difference in the outcomes between the two models. The THM variables correlated with each other and with the cleaning activities as it was also shown in the simple correlation plots. The correlations were less in number when the outcome was “any disease the past year” compared to the correlations visualized in the circo plot of the “general health” as an outcome. Additionally, different levels of the predictor variables were noted depending on the outcome, as it can be seen by the lines that are on the space outside the circular plot. These lines, however, were not used for interpretation due to the study’s exploratory nature to avoid any misleading inferences.

**Figure 5.**
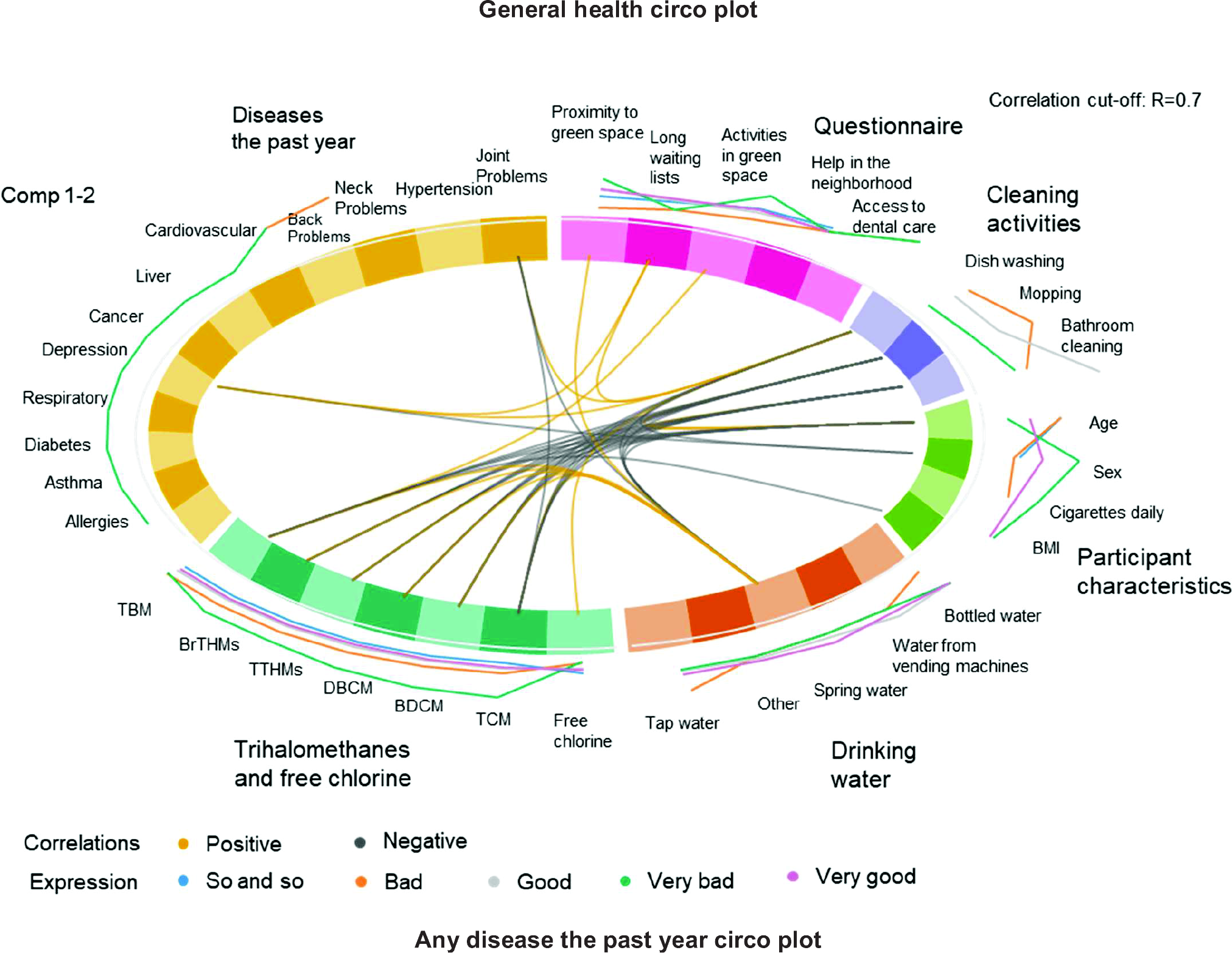
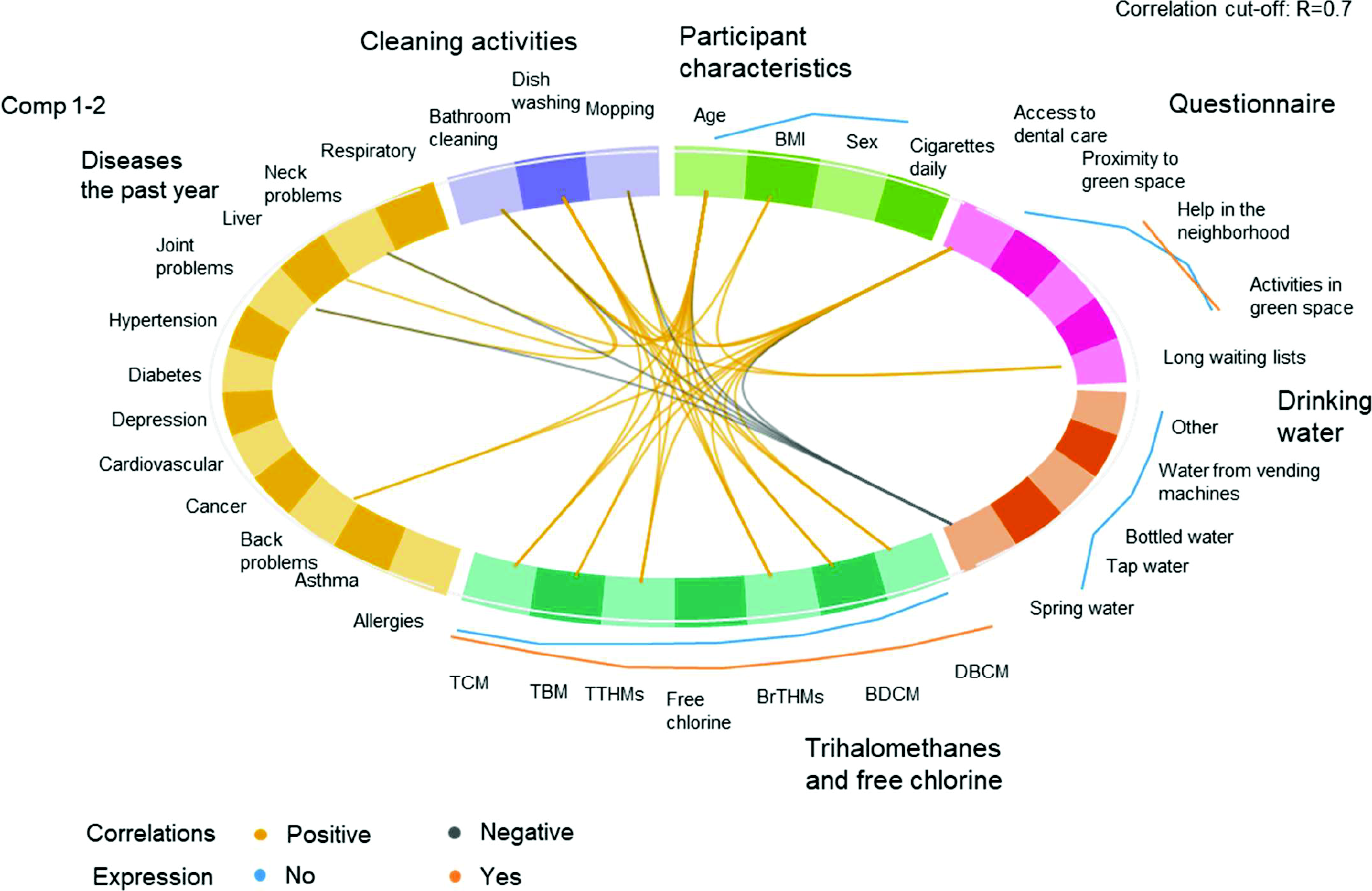
Circular plots of the correlations between the variables used in the environmental-wide analysis by block/group of variables.

## 4. Discussion

This is the first application, in a proof-of-concept study, of the urban exposome framework study in a medium-sized European city, Limassol, Cyprus that agnostically addressed and synthesized aspects of the life in the city (i.e. water quality and quality of life). The integrating nature of the urban exposome approach focused, in this instance, on the complexities of describing water quality indicators at the level of quarter, accounting for urban, general population characteristics, including the opinions/perceptions of residents and municipality stakeholders. The analysis included a suite of water quality aspects (chemical and microbial) and stakeholders’ opinions about environmental issues. Moreover, quality of life was assessed through citizens’ answers about access to healthcare and green spaces, and it was included along with lifestyle/behavior, and demographics in an EWAS analysis.

In this study, we used an interdisciplinary approach to identify trends in perceptions about environmental exposures and how they correlated with the actual water quality trends of the municipality of Limassol during the hot season (summer) with a combination of chemical and microbial indicators. Additionally, we generated global linkages and correlations between the health status of urban participants and their environmental/lifestyle/behavioral exposures at individual and quarter resolution levels. A total of 129 parameters from 36 variables either directly measured in water (water quality) or derived from the survey (quality of life in the city) were integrated using an exploratory, agnostic, EWAS approach. The general health and chronic health status of the urban participants were significantly associated in regression analysis with different health outcomes (e.g. hypertension, asthma) and quality of life indicators (e.g. financial issues in access to dental care). Circular correlation plots were derived from 36 urban exposome variables which were divided in six groups and accounting for self-reported health indicators (general health and having any diseases the past year). The correlation analysis revealed the association of water THM with cleaning activities, corroborating the overall results of simple correlations conducted within this study and our earlier results (24).

Our study did not reveal any unexpected results in terms of statistically significant associations between the variables included in this work. However, the combination of both the qualitative and quantitative parts allowed us to identify important community concerns about their urban life. Air pollution was ranked as the most significant concern among the study respondents. Besides the general interest of air pollution and its health effects, Cyprus experiences frequently dust storms (25). These events have probably triggered a specific concern among the population making air pollution the most frequently reported environmental concern in our study.

Besides a series of limitations such as the cross-sectional design, we were able to demonstrate the use of different tools in an integrative approach which aims to capture the urban exposome of Limassol in this case, but overall can be transferred to the study of the urban exposome of any city. The same approach, for example, if applied to larger cities will allow us to scale up the application of such integrative protocols and will allow for the wider transfer of the results. Future studies should, also, incorporate a more comprehensive assessment of urban quality of life (26). In this analysis, the indicators of quality of life were limited to life in the neighborhood or the use of green space. Additional information about social life is available from this study population and it will be incorporated in future studies. Characterising the urban exposome profile of Limassol will include from the routinely collected data of the registries and human biomonitoring analyses. Moreover, given the “stakeholders” assessment of the environmental problems, we have moved on with air quality measurements throughout the municipality which were conducted in spring 2018. This application shows how the goal of developing the urban exposome framework can be achieved by using all the available information in a real-time assessment of urban health and provide a tool for decision-making to stakeholders.

Urbanization, migration and other drivers of urban health are shaping the population health and quality of life in urban settings. Large scale neighborhood renewal programs are the so far initiatives to improve living conditions and quality of life in cities (27). The urban exposome as a study framework where cities and their smaller areas are the measurable units can be useful in monitoring urban health in a holistic manner. In other studies, the urban exposome defined as the totality of environmental exposures occurring in cities (28,29). So far, the use of human exposome in city-based cohorts has been focused on specific exposure-effect association testing in one or multiple cohorts being pooled together (28). However, the “urban exposome” vs. “specific health outcome/window of exposure” approach does not differ from the traditional environmental health and epidemiological study approaches and it carries the same limitations with regards to generalizability and high levels of variability among the studied populations or settings. In one of the first studies of this kind, where the urban exposome was studied with regards to its associations with pregnancy outcomes it was shown that extrapolating from multiple cities and from multiple populations comes with certain limitations which mostly pertain to the high between city variability (28). Thus, we need to move on and develop specific urban exposome studies where city-specific characteristics and within-city interactions and networks that are evidence-based, helping authorities to reach informed decisions about everyday life in the city, about city infrastructure changes and its effects on urban health and how it affects personal exposures. This will help the interpretation of between-city difference and will allow the timely evaluation of within-city challenges.

Our analysis did not raise any particular concerns about the quality of tap drinking water at the urban quarter scale of Limassol city using both chemical and microbial indicators. It was shown however, that residents do not trust, in general, the tap water and opt for using bottled water or water from other sources such as the “vending machine” water (groundwater from mountainous wells) which is very common practice in Cyprus. The cost of tap water is lower compared to bottled or “vending machine” water and, thus, it may pose additional burden to household budgets, as it has been shown in other studies (30). Mapping the cost of water by quarter would probably be informative about the existence or not of differences within Limassol in the economic burden of water consumption.

With the present proof-of-concept study we also mapped indicators of quality of life in the city including perceptions about access to green spaces, the life in the neighborhood and reasons for delay or financial constraint in access to health care. No clear disparities at the quarter level were observed for all neighborhood-based queries for the aforementioned indicators, but financial constraints for dental care were noted. Urban green space was particularly noted for being close to participants’ households, however, limited use of such green spaces (e.g. parks) was reported. Both aspects of access to health care and the use of green spaces within Limassol, were studied for the first time. Our approach could form the basis for future targeted and more integrative urban studies on these topics.

In general, our first results on the application of the urban exposome are promising and in need for verification and expansion. First and foremost, given the different habits of the citizens, exposome studies need to be more inclusive in the assessment of different water sources. Besides the inclusion of standard water testing parameters, future studies should address participant perceptions which are linked to behaviors, and, thus, exposures.

The case of the water presents one of the most challenging topics of urban setting to be studied within urban exposome studies due to the complex systems that drive water quality and water choices. These challenges are evident also in the literature about water quality indicators and their linkages with the exposome is overall scarce. For example, using an environment-wide association study methodology, the pregnancy exposome study on the INMA-Sabadell Birth Cohort in Spain looked into water disinfection byproducts (trihalomethanes) modeled previously at the individual level among a wider set of other exposures ranging from urinary markers of chemical exposures to air pollution and noise (3); they found that the three trihalomethane “classes” studied (total trihalomethanes, brominated trihalomethanes and chloroform) were strongly correlated with each other, but not with the other exposures (3). Albouy-Llaty et al (2016), explored the association between endocrine disruptors (atrazine metabolites and nitrate/atrazine mixture) in drinking water and preterm birth accounting for socioeconomic factors (deprivation index) in the Poitou-Charentes region of France (31). The exposure to atrazine and the nitrate/atrazine mixture at the individual level was inferred from routine community monitoring of water quality; preterm birth was found to be associated with the deprivation index at the level of the neighborhood but not with the exposure to atrazine metabolites. Exposure to atrazine (measured through the metabolite 2-hydroxyatrazine) was not found to be a significant risk factor for preterm birth when accounting for the socioeconomic status of the area (31).

Developing the health profile of a city in urban exposome terms and integrating different approaches comes with several limitations. Spatio-temporal considerations should be accounted for the dynamic nature of urban exposome profiling. Although enough to estimate the background levels of population exposures to chemicals, the relatively small sample size might have not allowed us to capture spatial differences of the indicators measured within the smaller city administrative units. Additionally, the lack of biomonitoring data on water-related exposures (i.e. to disinfection byproducts in urine) have hindered the full deployment of EWAS tool capabilities. However, the availability of urine biospecimen for this survey will allow us to use biomonitoring and untargeted metabolomics tools in a follow-up manuscript.

## 5. Conclusions

Developing city health profiles with the aid of the urban exposome framework is a novel approach, yet, far from being simple. It demands a comprehensive characterization of relevant indicators ranging from drinking water quality to health perceptions and opinions, etc. The urban exposome framework and its application will generate tools and datasets that will pave the way for developing the next innovative solutions and public health interventions for the city. This case study of the urban exposome in Limassol, Cyprus, demonstrates the feasibility of using novel exposome approaches in studying the city and smaller within-city areas (quarters) as the units of reference. Within this context, the absolute water quality indicators, city residents’ and other stakeholders’ opinions need to be integrated and expanded along with exposomic profiles, such as metabolomics and other-omics platforms and human biomonitoring protocols.

## Author Contributions

Original idea: X.D.A. and K.C.M. Citizens perceptions study design and analysis: C.vdL., X.D.A, and C.B. Participant recruitment X.D.A, C.vdL, S.I.; Water sample analysis for disinfection by-products: P.C. and K.F.; Free chlorine measurements: S.I.; Microbiological analysis: Z.P and G.B.; Data analysis for the population study: X.D.A; Manuscript preparation: X.D.A. and K.C.M. All authors reviewed and provided feedback on earlier versions of the manuscript.

## Acknowledgments

We would like to thank the Municipality of Limassol and all the study participants. Special thanks to Ms. Andriana Till for her contribution to participant recruitment and to Dr. Stephanie Gaengler for the fruitful discussions during data analysis. We would also like to express our gratitude to Drs. Athos Agapiou and Diofantos Hadjimitsis for sharing the map templates for the quarters of Limassol.

