## Supplemental material for "Mapping drinking water and quality of life aspects in urban settings: an application of the urban exposome"

The supplementary information includes additional tables and figures and the methodology of the microbiological analysis.

### Additional tables and figures

**Table S 1 Population by quarter, sample size estimation and number of participants.**

| | GEO CODE* | Area | Total (Census 2011**) | Estimated sample size | Recruited |
|---|---|---|---|---|---|
| Municipality | 5000 | Limassol | 101000 | 120 | 132 |
| Quarters | 500017 | Agios Nikolaos | 5631 | 7 | 5 |
| | 500013 | Agios Nektarios | 3397 | 4 | 3 |
| | 500012 | Kapsalos | 6660 | 8 | 7 |
| | 500015 | Agia Trias | 2786 | 3 | 3 |
| | 500016 | Neapoli | 7229 | 9 | 10 |
| | 500008 | Omonoia | 3839 | 5 | 5 |
| | 500014 | Agia Zoni | 4456 | 5 | 5 |
| | 500020 | Agios Spyridon | 9439 | 11 | 11 |
| | 500021 | Zakaki | 5874 | 7 | 8 |
| | 500011 | Apostoloi Petros kai Pavlos | 10412 | 12 | 14 |
| | 500009 | Apostolos Andreas | 9207 | 11 | 14 |
| | 500010 | Agios Georgios | 5060 | 6 | 9 |
| | 500005 | Katholiki | 4647 | 6 | 12 |
| Combined quarters (presented separately and summed together) | 500004 | Agia Napa | 534 | 1 | 1 |
| | 500002 | Tziami Tzentit | 434 | 1 | 1 |
| | 500007 | Tsiflikoudia | 579 | 1 | 1 |
| | | (Beachfront quarters) | 1547 | 3 | 3 |
| | 500018 | Agia Fylaxis | 14451 | 17 | 17 |
| | 500019 | Panagia Evangelistria | 693 | 1 | 1 |
| | | (Agia Fylaxis and Panagia Evangelistria) | 15144 | 18 | 18 |
| | 500003 | Arnaoutogeitonia | 905 | 1 | 1 |
| | 500006 | Agios Ioannis | 4767 | 6 | 4 |
| | | (Arnaoutogeitonia and Agios Ioannis) | 5672 | 7 | 5 |

*Based ont he 2011 Population Census of Cyprus

**Source: Population Census 2011 [Internet]. Statistical Service, Republic of Cyprus. 2014 [cited 2014 Sep 9]. Available from: http://www.mof.gov.cy/mof/cystat/statistics.nsf/census-2011_cystat_en/census-2011_cystat_en?OpenDocument

**Table S 2 Variables from the urban population study questionnaire used in the environment-wide association study (EWAS) analysis. Part A of the table lists the outcomes, and Part B the predictors by block/group.**

| **Part A** | **Outcomes** | | |
|---|---|---|---|
| **Comment on the predictors** | **Variable** | **Categories( if applicable)** | **Score** |
| Predictors from blocks/groups 1-6 (number of predictor variables=36) | General health | Very good | 2 |
| | | Good | 1 |
| | | So and so | 0 |
| | | Bad | -1 |
| | | Very bad | -2 |
| | Chronic disease | Yes | 1 |
| | | No | 0 |
| | | I don't know/I don't want to answer | NA |
| Predictors from blocks/groups 1-5 (number of predictor variables=25) | Any disease past year | Yes | 1 |
| | | No | 0 |
| **Part B** | **Predictors** | | |
| **Block/Group (number of variables)** | **Description of variables** | **Categories (if applicable)** | **Score** |
| (B1) Chemical water indicators (n=7) | Trihalomethanes (Total THM, BrTHM, TCM, BDCM, DBCM, TBM; units: µg/L)<br>Free chlorine (units mg/L; ND=0) | | |
| (B2) Drinking water habits (n=5) | Number of glasses of water per day by source | | |
| (B3) Cleaning activities (n=3) | Mopping, bathroom cleaning, dishwashing (times per week) | | |
| (B4) Questionnaire variables (n=5) | | I don't want to answer | NA |
| | Delays in access to health care services due to long waiting lists | No, I didn't face any delays (reference) | +1 |
| | | No, I didn't need care | 0 |
| | | Yes | -1 |
| | | I don't want to answer | NA |
| | Financial constraints in access to dental care | No, I could afford it (reference) | +1 |
| | | No, I didn't need | 0 |
| | | Yes | -1 |
| | Living close to green space (proximity)<br>Can do many activities in the green space nearby<br>There is always someone to help you in the neighborhood | Completely agree (reference) | +2 |
| | | Probably agree | +1 |
| | | Do not know | 0 |
| | | Probably disagree | -1 |
| | | Completely disagree | -2 |
| (B5) Participant characteristics (n=5) | Age (years)<br>BMI (kg/m^2)<br>Number of cigarettes smoking daily | | |
| | Sex | Female (reference) | 0 |
| | | Male | 1 |
| (B6) Self-reported diseases the past year (n=11) | Asthma, respiratory diseases, hypertension, cardiovascular diseases, joint or other muscloskeletal problems, diabetes, allergies, liver disorders, cancer, depression | Yes | 1 |
| | | No (reference) | 0 |
| | | I don't know/<br>I don't want to answer | NA |

Notes: NA: excluded as missing

**Table S 3 Additional characteristics of the study population based on the questionnaire responses of the urban population study.**

| | Overall (n=132) |
|---|---|
| **Years living in Cyprus (%)** | |
| Less than a year | 3 ( 2.3) |
| 1-5 years | 6 ( 4.5) |
| 6-10 years | 4 ( 3.0) |
| 11-20 years | 3 ( 2.3) |
| More than 20 years | 20 (15.2) |
| All my life | 96 (72.7) |
| **Place of birth (%)** | |
| Cyprus | 114 (86.4) |
| Other EU country | 13 ( 9.8) |
| Other non-EU country | 5 ( 3.8) |
| Smoking status (%) | |
| I don't want to answer | 1 ( 0.8) |
| Non-smokers | 81 (61.8) |
| Smokers | 41 (31.3) |
| Occasional smoking | 8 ( 6.1) |
| **Exposure to secondhand smoke (hours per day) (%)** | |
| I don't know/I don't remember | 8 ( 6.2) |
| I don't want to answer | 1 ( 0.8) |
| Less than 1 hour per day | 35 (27.3) |
| More than 1 hour per day | 32 (25.0) |
| Never or almost never | 52 (40.6) |
| **Alcoholic consumption the past 12 months (%)** | |
| Daily or almost daily | 11 ( 8.4) |
| 5-6 times per week | 2 ( 1.5) |
| 3-4 times per week | 15 (11.5) |
| 1-2 times per week | 33 (25.2) |
| 2-3 days per month | 23 (17.6) |
| 1 time per month | 8 ( 6.1) |
| Less than 1 time per month | 15 (11.5) |
| Never consumed alcohol in the past 12 months | 7 (5.3) |
| Never or I have consumed alcohol only a few times in my life | 13 (9.9) |
| I don't want to answer | 4 ( 3.1) |

**Table S 4 Summary of the responses to different questions relating to health care access, lifestyle and quality of life in the neighborhood.**

| | | Overall (n=132) |
|---|---|---|
| **Delays in access to health care services** | | |
| Delays in health care due to long waiting list (%) | I don't want to answer | 6 (4.5) |
| | No, I didn't face any delays | 43 (32.6) |
| | No, I didn't need care | 68 (51.5) |
| | Yes | 15 (11.4) |
| Delays in health care due to lack of transport (%) | I don't want to answer | 20 (15.2) |
| | No, I didn't face any delays | 43 (32.6) |
| | No, I didn't need care | 67 (50.8) |
| | Yes | 2 (1.5) |
| Financial constraints in access to medical care (%) | I don't want to answer | 9 (6.8) |
| | No, I could afford it | 45 (34.1) |
| | No, I didn't need | 75 (56.8) |
| | Yes | 3 (2.3) |
| **Financial constraints** | | |
| Financial constraints in access to dental care (%) | I don't want to answer | 4 (3.0) |
| | No, I could afford it | 40 (30.3) |
| | No, I didn't need | 70 (53.0) |
| | Yes | 18 (13.6) |
| Financial constraints in access to buy any medications (%) | I don't want to answer | 6 (4.5) |
| | No, I could afford it | 45 (34.1) |
| | No, I didn't need | 76 (57.6) |
| | Yes | 5 (3.8) |
| Financial constraints in access to mental health care (%) | I don't want to answer | 11 (8.3) |
| | No, I could afford it | 5 (3.8) |
| | No, I didn't need | 113 (85.6) |
| | Yes | 3 (2.3) |
| **Opinions about green space near the residence** | | |
| Enough green spaces (%) | Completely agree | 18 (13.6) |
| | Probably agree | 29 (22.0) |
| | Do not know | 8 (6.1) |
| | Probably disagree | 45 (34.1) |
| | Completely disagree | 32 (24.2) |
| Access to green spaces is easy (%) | Completely agree | 38 (28.8) |
| | Probably agree | 39 (29.5) |
| | Do not know | 9 ( 6.8) |
| | Probably disagree | 24 (18.2) |
| | Completely disagree | 22 (16.7) |
| Living close to green space (proximity) (%) | Completely agree | 45 (34.1) |
| | Probably agree | 40 (30.3) |
| | Do not know | 7 ( 5.3) |
| | Probably disagree | 17 (12.9) |
| | Completely disagree | 23 (17.4) |
| Green spaces nearby are well-maintained (%) | Completely agree | 11 ( 8.3) |
| | Probably agree | 39 (29.5) |
| | Do not know | 12 ( 9.1) |

| | | |
|---|---|---|
| | Probably disagree | 39 (29.5) |
| | Completely disagree | 31 (23.5) |
| Relaxing in the green spaces nearby (%) | Completely agree | 14 (10.6) |
| | Probably agree | 26 (19.7) |
| | Do not know | 6 ( 4.5) |
| | Probably disagree | 39 (29.5) |
| | Completely disagree | 47 (35.6) |
| Can do many activities in green space (%) | Completely agree | 8 ( 6.1) |
| | Probably agree | 18 (13.6) |
| | Do not know | 9 ( 6.8) |
| | Probably disagree | 46 (34.8) |
| | Completely disagree | 51 (38.6) |
| **Opinions about different aspects of life in the neighborhood** | | |
| The neighbors are willing to help each other (%) | Completely agree | 41 (31.1) |
| | Probably agree | 59 (44.7) |
| | Do not know | 13 ( 9.8) |
| | Probably disagree | 14 (10.6) |
| | Completely disagree | 5 ( 3.8) |
| Neighbors share the same values (%) | Completely agree | 34 (25.8) |
| | Probably agree | 57 (43.2) |
| | Do not know | 23 (17.4) |
| | Probably disagree | 12 ( 9.1) |
| | Completely disagree | 6 ( 4.5) |
| There is always someone to ask help you(%) | Completely agree | 51 (38.6) |
| | Probably agree | 56 (42.4) |
| | Do not know | 15 (11.4) |
| | Probably disagree | 6 ( 4.5) |
| | Completely disagree | 4 ( 3.0) |

**Table S 5 The first 20 parameters from the univariate models ranked by FDR adjusted p-value. In the categorical outcomes (i.e. chronic disease and any disease the past year, noted as "ChronicDisease" and "Disease12M" in the column "Outcome") the estimate is the odds ratio**.

| Variable | Term | Estimate | p value | 5% CI | 95% CI | n (%) | Outcome | FDR |
|---|---|---|---|---|---|---|---|---|
| FinancialIssuesDentalCareREC | valueYes | -0.794 | 0 | -1.167 | -0.422 | 18 (14.1) | GeneralHealth | 0 |
| Depression12MREC | valueYes | -1.448 | 0 | -2.195 | -0.7 | 3 (2.8) | GeneralHealth | 0 |
| Hypertension12MREC | valueYes | 11.714 | 0 | 3.858 | 38.429 | 19 (16.2) | ChronicDisease | 0 |
| Age | value | 3.293 | 0 | 2.128 | 5.337 | NA | Disease12M | 0 |
| JointProblems12MREC | valueYes | -0.621 | 0.001 | -0.998 | -0.244 | 12 (10.8) | GeneralHealth | 0.0215 |
| BackProblems12MREC | valueYes | -0.5 | 0.001 | -0.782 | -0.218 | 37 (32.2) | GeneralHealth | 0.0215 |
| Age | value | -0.194 | 0.002 | -0.318 | -0.07 | NA | GeneralHealth | 0.036857 |
| NeckProblems12MREC | valueYes | -0.479 | 0.003 | -0.79 | -0.168 | 20 (17.5) | GeneralHealth | 0.043 |
| BackProblems12MREC | valueYes | 4.402 | 0.003 | 1.67 | 12.124 | 37 (32.2) | ChronicDisease | 0.043 |
| Age | value | 2.074 | 0.004 | 1.297 | 3.589 | NA | ChronicDisease | 0.046909 |
| NeckProblems12MREC | valueYes | 4.657 | 0.004 | 1.602 | 13.623 | 20 (17.5) | ChronicDisease | 0.046909 |
| Asthma12MREC | valueYes | 6.071 | 0.009 | 1.515 | 24.61 | 10 (8.5) | ChronicDisease | 0.09675 |
| ProximityGreenSpace | valueDo not know | -0.708 | 0.018 | -1.292 | -0.124 | 7 (5.3) | GeneralHealth | 0.175071 |
| LongWaitingListHCDelay12MREC | valueYes | 4.5 | 0.019 | 1.311 | 16.672 | 15 (11.9) | ChronicDisease | 0.175071 |
| Hypertension12MREC | valueYes | -0.354 | 0.024 | -0.661 | -0.046 | 19 (16.2) | GeneralHealth | 0.2064 |
| Alergies12MREC | valueYes | 3.846 | 0.026 | 1.129 | 12.624 | 17 (15.5) | ChronicDisease | 0.209625 |
| Mopping_days_week | value | -0.135 | 0.035 | -0.261 | -0.01 | NA | GeneralHealth | 0.265588 |
| ActivitiesInGreenSpace | valueDo not know | -0.736 | 0.04 | -1.437 | -0.035 | 9 (6.8) | GeneralHealth | 0.278368 |
| Spring_water_glasses_day | value | 0.506 | 0.041 | 0.191 | 0.835 | NA | Disease12M | 0.278368 |
| Dish_washing_days_week | value | 1.431 | | 1.009 | 2.058 | NA | Disease12M | 0.30315 |

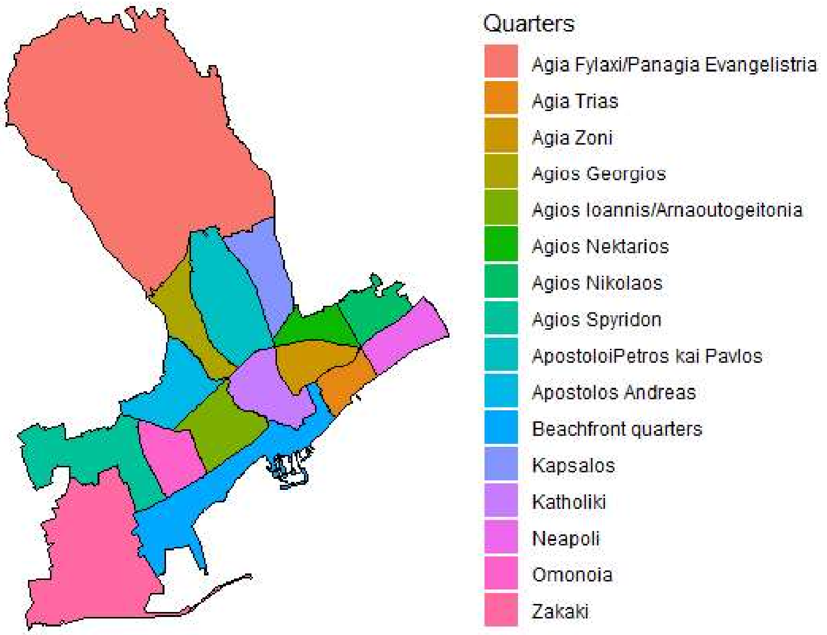


**Figure S 1 Map of the quarters of the Limassol municipality as they are used in the analysis.**

**R packages used in the data analysis.**

| Packages used in the analysis |
|---|
| [1] mixOmics_6.3.2 MASS_7.3-50 gdtools_0.1.7 |
| [4] bindrcpp_0.2.2 rJava_0.9-10 xlsx_0.6.1 |
| [7] officer_0.3.2 rvg_0.1.9 scales_0.5.0 |
| [10] viridis_0.5.1 viridisLite_0.3.0 broom_0.4.5 |
| [13] reshape2_1.4.3 knitr_1.20 tableone_0.9.3 |
| [16] ISOweek_0.6-2 summarytools_0.8.5 corrplot_0.84 |
| [19] Hmisc_4.1-1 Formula_1.2-3 survival_2.42-3 |
| [22] lattice_0.20-35 forcats_0.3.0 stringr_1.3.1 |
| [25] purrr_0.2.5 readr_1.1.1 tidyr_0.8.1 |
| [28] tibble_1.4.2 tidyverse_1.2.1 lubridate_1.7.4 |
| [31] readxl_1.1.0 RColorBrewer_1.1-2 dplyr_0.7.6 |
| [34] data.table_1.11.4 rgdal_1.3-3 sp_1.3-1 |
| [37] plyr_1.8.4 ggplot2_3.0.0 |

### Microbiological analysis

**Total viable counts at 22 and 37 °C**

Enumeration of heterotrophic bacteria in water was performed by the pour plate method with yeast extract agar (YEA; Oxoid, CM0019, Basingstoke, UK) (Suthar et al. 2009). Briefly, 1 ml of water sample was transferred onto sterile 90-mm Petri dishes, followed by the addition of 15 ml of YEA (previously autoclaved and cooled to 45–50 °C). The contents were mixed by a combination of rapid end-to end shaking and circular movements lasting over a period of 5–10 s. The agar was then allowed to solidify, and incubation of duplicate sets of plates for 3 days at 37 and at 22 °C was practiced.

**Membrane filtration analysis for *E. coli*, coliforms, *Pseudomonas aeruginosa*, and *Enterococus* spp.**

The membrane filtration technique was applied for the detection and enumeration of *E. coli*, coliforms, *Pseudomonas aeruginosa*, and *Enterococcus* spp., followed by incubation onto selective media. A volume of 250 ml of the sample was filtered through a 0.45-mm, gridded, sterile membrane, and the filter was then aseptically transferred onto the appropriate agar medium in Petri dish, avoiding air bubbles beneath the membrane. For the *E. coli* and coliform analysis, a cromogenic medium was used (ChromoCult® Coliform Agar, Merck, Darmstadt, Germany), which was able to differentiate between *E. coli* and other coliform bacteria. Plates were incubated inverted at 37 °C for 24 h with red color colonies, indicating the presence of coliform bacteria, whereas the presence of blue colonies indicated the presence of *E. coli* (see Ouattara et al. 2011).

The analysis for *Pseudomonas aeruginosa* was performed onto Pseudomonas Agar (OXOID CM0559, plus Pseudomonas CN selective agar supplement SR0102, Basingstoke, UK). Incubation was set at 25 °C for 2 days with positive colonies of *Pseudomonas aeruginosa* coming up with light green color and fluoresced under UV light at 365 nm. Finally, for enumerating enterococci from the water samples, the membrane filters were incubated onto Slanetz and Bartley medium (OXOID CM0377, Basingstoke, UK) at 44 °C for 4 h and at 37 °C for 2 days. Colonies with red-brown color were enumerated as *Enterococcus* spp. All suspicious colonies were confirmed molecularly via 16S rRNA sequencing as previously described by Botsaris et al. (2015).
